# Secretion and Periplasmic Activation of a Potent Endonuclease in E. coli

**DOI:** 10.1101/2024.04.16.589800

**Authors:** Mehran Soltani, James R. Swartz

## Abstract

Sm Endonuclease (SmEn) is a promiscuous, highly active nuclease widely used in protein purification, 2D protein gels, and gene and cell therapy. We aimed to recombinantly and economically produce this reagent using E. coli. Despite widespread application of E. coli for recombinant production of proteins, cytoplasmic expression of this protein resulted in no activity accumulation. We therefore investigated translocation of SmEn to the periplasm of E. coli by evaluating several signal sequences, E. coli host cells, and incubation conditions. For rapid feedback, we developed a crude lysate-based nuclease activity assay that enabled convenient screening and identified suitable conditions for active SmEn accumulation. Signal sequence selection was most influential with additional benefit gained by slowing synthesis either using the transcriptionally weakened strain, C43 (DE3) or by reducing incubation temperature. While our study provides valuable insights for optimizing a nuclease translocation and reducing production costs, more research is needed to explore the influence of mRNA secondary structure at the translation initiation region on protein expression and translocation. Overall, our rapid screening assay facilitated the development of an effective production process for a protein with potential cytoplasmic toxicity as well as the need of disulfide bond formation.

## Introduction

Sm Endonuclease (SmEn; endonuclease from Serratia marcescens, PDB code: 1G8T) is a non-specific nuclease that digests double stranded and single stranded DNAs and RNAs to 2-5 base pair oligonucleotides [1]. SmEn facilitates cell lysate handling and protein purification by digesting genomic DNA to reduce lysate viscosity [2, 3]. It is also used to minimize nucleic acids carry-over for immunoblotting [4, 5], western blotting [3, 6], purifying viral vectors used for gene therapy, and sample preparation for 2D protein gels [7, 8]. Even though this reagent is commercially available from Millipore (as Benzonase^®^) and c-LECTA (as DENARASE^®^), it is expensive (more than $6000 per mg from Millipore) especially for large-scale lysate preparation.

Here, we describe the development of a cost-effective SmEn production process. E. coli was selected as the production host due to its rapid growth rate, high heterogeneous protein yields, simple cultivation, and inexpensive growth media [9]. Considering the potential toxicity of cytoplasmically accumulated SmEn and the requirement to form two disulfide bonds for activity and stability [10], we aimed to translocate SmEn to the periplasm of E. coli, where DsbA and DsbC could assist oxidative protein folding [11]. Secretion of proteins into the periplasmic space followed by osmotic extraction also potentially reduces contamination of product proteins with endotoxins and native E. coli proteins to reduce downstream processing costs compared to cytoplasmic expression [12]. Periplasmic accumulated may also improve folding because of lower protein-protein interactions compared to cytoplasmic expression, and the addition of an N-terminal methionine is avoided [13, 14].

For protein translocation across the cytoplasmic membrane, six secretion signal sequences (signal peptides) were tested to evaluate both Sec and Tat (Twin-arginine-translocation) transport processes. The Sec secretion sequences of PhoA, Hbp, ST2, DsbA, and OmpA were evaluated. Considering the possibility of SmEn cytoplasmic toxicity because of the translocation of fully folded proteins by the Tat pathway, we only tested one Tat signal sequence (TorA or Tri-methylamine-N-oxide reductase) [15].

Sec-translocation is the dominant translocation pathway in E. coli [16]. This mechanism is believed to either co- or post-translationally translocate SecB-bound or SRP-directed polypeptides targeted for translocation by an N-terminal signal sequence [17-20]. This production mode is expected to avoid possible cytoplasmic SmEn toxicity and periplasmic protein folding can be facilitated after signal peptide removal by signal peptidase by the disulfide bond forming and isomerizing oxidoreductases DsbA and DsbC) [21, 22].

In addition to the commonly used E. coli BL21 (DE3) strain, we also evaluated a C43 (DE3) strain known as the Walker strain often used to provide higher membrane and secreted protein accumulation. This strain has multiple mutations that weaken the lacUV5 promoter used for T7 RNA polymerase expression [21, 23, 24], thereby slowing transcription initiation rates for target gene expression. Slower production can avoid saturating translocation capacity which otherwise could lead to inclusion body formation, translocon inhibition, and/or growth arrest [25-28].

To evaluate the effect of protein synthesis rate, we tested various incubation temperatures and induction levels. A new convenient screening assay directly assessed nuclease activity reducing assessment time to less than 1.5 hours and eliminating the need for laborious cell lysis, purification, and buffer exchange. Additionally, it was important to consider several previous reports of soluble but misfolded and inactive secreted protein accumulation [29-31] which indicated the need to optimize SmEn translocation and folding based on nuclease activity and not protein accumulation.

## Materials and Methods

### Plasmid preparation and RNA secondary structure determination

The amino acid sequence for SmEn without a signal sequence was obtained from the Protein Data Bank (PDB: 1G8T). The DNA sequence was codon-optimized for E. coli translation and constructed by Twist Bioscience (South San Francisco, CA). The gene was cloned into pET21b (Novagen) via Gibson assembly and extended to encode a C-terminal 6x Histidine tag to facilitate affinity purification. DNA sequences coding for six different signal peptides were constructed by Twist Bioscience (DNA sequences in the Supplementary Information), inserted at the N-terminus of the SmEn gene using Gibson assembly, and verified by DNA sequencing. DNA sequences for these signal sequences were obtained from previously published works [24, 32]. Assembled SmEn constructs with different signal peptides in the pET21b backbone were transformed into BL21 (DE3) (Invitrogen) or OverExpress™ C43 (DE3) (Sigma-Aldrich) chemically competent cells.

To assess possible RNA secondary structures, the nucleotide sequence of the 5’ untranslated region (UTR), ribosome binding site region (30 bps) and the initial 40 bps of the gene (variable for different signal sequences) were analyzed using default parameters by RNAstructure Fold, an online web server developed by Mathews et al. [33].

### Cell culture

All cell cultures were performed as described previously [34] with slight modification. Briefly, 3 ml LB media containing 100 µg/ml carbenicillin was inoculated and incubated overnight and with shaking at 250 RPM at 37 °C. The cells were then transferred to 250 ml fresh 2x YTPG media (16 g/L tryptone, 10 g/L yeast extract, 7 g/L potassium phosphate dibasic, 3 g/L potassium phosphate monobasic, 18 g/L glucose, pH adjusted to 7.2 by KOH) [35] in 500 ml baffled shake flasks (Greiner Bio-One). Incubation was conducted at 37 °C and 250 RPM agitation until reaching an OD600 of ∼0.5-0.6. Cultures were then induced by 10-250 µM isopropyl β-D-thiogalactoside (IPTG). Incubation continued at different temperatures (15-37 °C) overnight (∼15 h) with a final OD600 of 3-3.8 depending on culture condition and signal peptide. Cells were harvested at 8000 g for 20 min at 4 °C and stored at -80 °C until evaluation.

### SmEn isolation and purification

To recover SmEn for the activity assay, 0.5 ±0.001 gr wet cells was resuspended in 2.25 ml buffer Z (40 mM Tris, 20 mM NaCl, 2 mM MgSO4, pH=8.0), and 0.25 ml 10 X endonuclease-free BugBuster Protein Extraction Reagent (Millipore, Cat. No. 70921) was added. The mixture then was incubated at 37 °C with stirring for 30 min before centrifugation at 15000 g for 20 min at 4 °C. The supernatant was aliquoted and tested (freshly without freeze-thaw cycles) for nuclease activity.

For SmEn affinity purification, cells with translocated SmEn were harvested, resuspended in buffer Z (2 ml/g wet cells), lysed by a single pass through an EmulsiFlex-C5 homogenizer (Avestin inc., Ottawa, Canada), and centrifugated at 15000 g for 20 min at 4 °C. Supernatant lysate was collected and SmEn was purified in the gravity flow mode using Ni-NTA resin (G-bioscience, Cat. # 786-1547) following the manufacturer’s recommended procedure with slight modifications. Briefly, 1 ml Ni-NTA resin was used per gram wet cells resuspended in 2 ml Z buffer (supplemented with 20 mM imidazole) and the same buffer was used as the wash buffer (5 column volumes). Buffer Z supplemented with 50-300 mM imidazole was used to elute SmEn in a step-gradient mode (each step was 0.5 column volume) as indicated.

For protein visualization, 4-12% NuPAGE™ gels were used per manufacture’s recommended protocol with MES as the running buffer. Cell lysate before loading to N-NTA resin, flow through, and wash solutions were all diluted 15 times before loading onto the protein gel. Elution fractions were loaded without dilution. The DC protein assay (Bio-Rad, Hercules, CA) was used for protein concentration measurement according to the manufacturer’s protocol using bovine serum albumin as the calibration standard.

### Nuclease activity assay

The nuclease activity assay used 0.74 µg/µl salmon sperm DNA (Invitrogen, Cat. No. 15632011) as the DNA substrate and variable dilutions of lysate (0.005 to 5% (v/v)) in buffer Z to a 50 µl final volume. Reaction mixtures were incubated at 37 °C for 30 min and then maintained at 4 °C to stop the DNA degradation reaction until dilution with gel loading dye containing 10 mM EDTA to stop the DNA degradation reaction. To visualize residual DNA from the nuclease assay reactions, 7.2 µl of the assay reaction mixture was added to 3 µl of 6X Gel Loading Dye, Purple (NEB, Cat. No. B7024) and 7.8 µl MilliQ water (18 µl final volume). 14 µl of the resulting mixture was loaded on a 1% agarose gel and electrophoresed at 110 V for 15 min, followed by DNA visualization by UV light. As a negative control, Z buffer was used instead of SmEn lysates.

## Results and Discussion

### Screening assay for suitable signal peptide and E. coli host

First, we developed a novel and rapid screening assay to assess periplasmic SmEn activity accumulation. This assay visualizes the digestion of salmon sperm DNA by different dilutions of crude lysates prepared from E. coli cells secreting SmEn, thereby indicating relative SmEn activity accumulation. The results shown in Figure 1 indicate typical assay performance.

**Figure 1.**
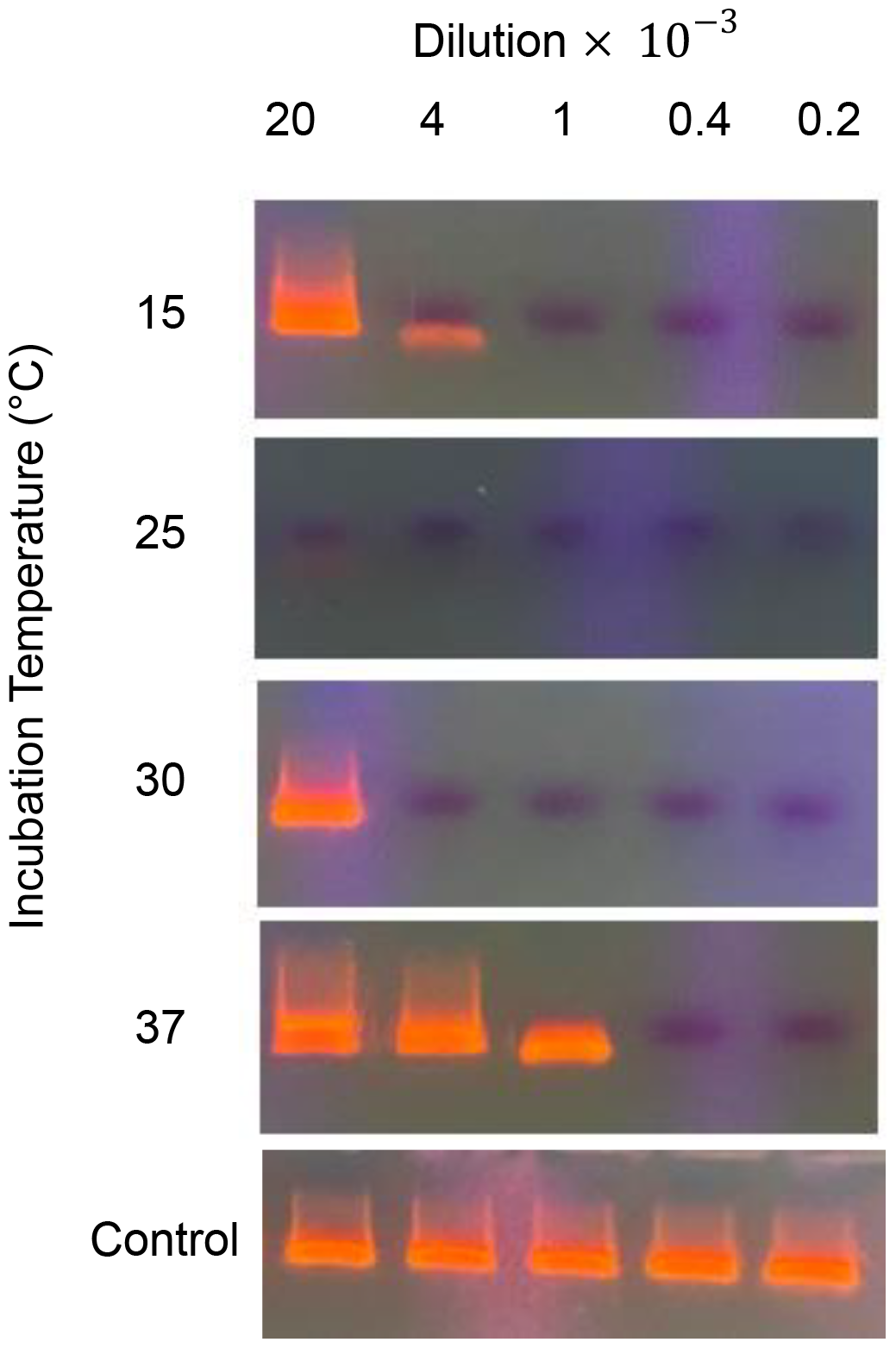
Incubation temperature effect on SmEn activity production using the Hbp signal peptide in C43 (DE3).

IPTG titration using the C43 (DE3) host and the Hbp signal peptide was first conducted to provide a baseline activity standard. The same accumulation of SmEn activity in the periplasm was observed regardless of IPTG concentration for C43 (DE3) cells at 30 °C (Figure S1). Inability to modify T7 RNA polymerase expression for the control of transcription rate using different IPTG concentrations (approximating constitutive expression) has been discussed previously and attributed to active IPTG uptake by cells either using Lac permease (lacY) or by permease independent pathways [9, 36].

Consequently, different post-induction incubation temperatures (15, 25, 30, and 37 °C) were investigated to improve periplasmic SmEn activity accumulation using the Hbp signal peptide in the C43 (DE3) host. All cells were induced by 150 µM IPTG at an OD600 of 0.5-0.6 and incubated overnight at the various temperatures. Figure 1 shows SmEn activity indicated by the 30-minute DNA degradation activity assay described in Materials and Methods. Room temperature incubation resulted in the highest SmEn activity as indicated by complete DNA band disappearance at the highest dilution (20,000 fold).

Improved SmEn activities after room temperature incubation could be attributed to a slower protein synthesis rate at the lower temperature to facilitate more effective translocation across the inner membrane. Higher temperatures that promote rapid transcription and translation can overwhelm the translocation machinery to result in toxic accumulation of target protein aggregates that inhibit the translocon or other essential functions [27, 37]. Lower incubation temperature could also avoid overwhelming DsbA and DsbC’s capacity for aiding oxidative protein folding.

The need for slower translation and translocation rates for optimal accumulation of properly folded secreted heterologous proteins has been demonstrated previously by varying the nucleotide sequence of the translation initiation region (TIRs, codons 2 to 6) without altering the amino acid sequence while using the Heat-Stable Enterotoxin II signal peptide (ST2) [32]. Slower translation initiation rates improved human protein secretion in all four cases evaluated.

Based on the preliminary results, six different signal peptides and two E. coli host strains were tested for SmEn production with 150 µM IPTG induction at an OD600 of about 0.5 and overnight incubation at room temperature. Final OD600’s were typically somewhat lower for the C43 (DE3) host (Table S1) with an average of 4.10 for BL21 (DE3) and 3.46 for C43 (DE3).

Figure 2 shows the resulting activity assessments. As expected, the TorA signal peptide that directs secretion through the Tat pathway resulted in the lowest accumulation of SmEn activity. Even though the protein could become fully folded before translocation by this mechanism [38, 39], no apparent cytoplasmic toxicity was indicated by final cell densities. The highest nuclease activity was produced using the Hbp signal peptide in C43 (DE3) host cells followed by the OmpA signal peptide in either BL21 (DE3) or C43 (DE3) host cells. Lower SmEn accumulation with BL21 (DE3) compared to C43 (DE3) can be attributed to faster SmEn mRNA production in BL21 (DE3) [40] which could overwhelm the translocation apparatus to then inhibit translation, translocation, or both.

**Figure 2.**
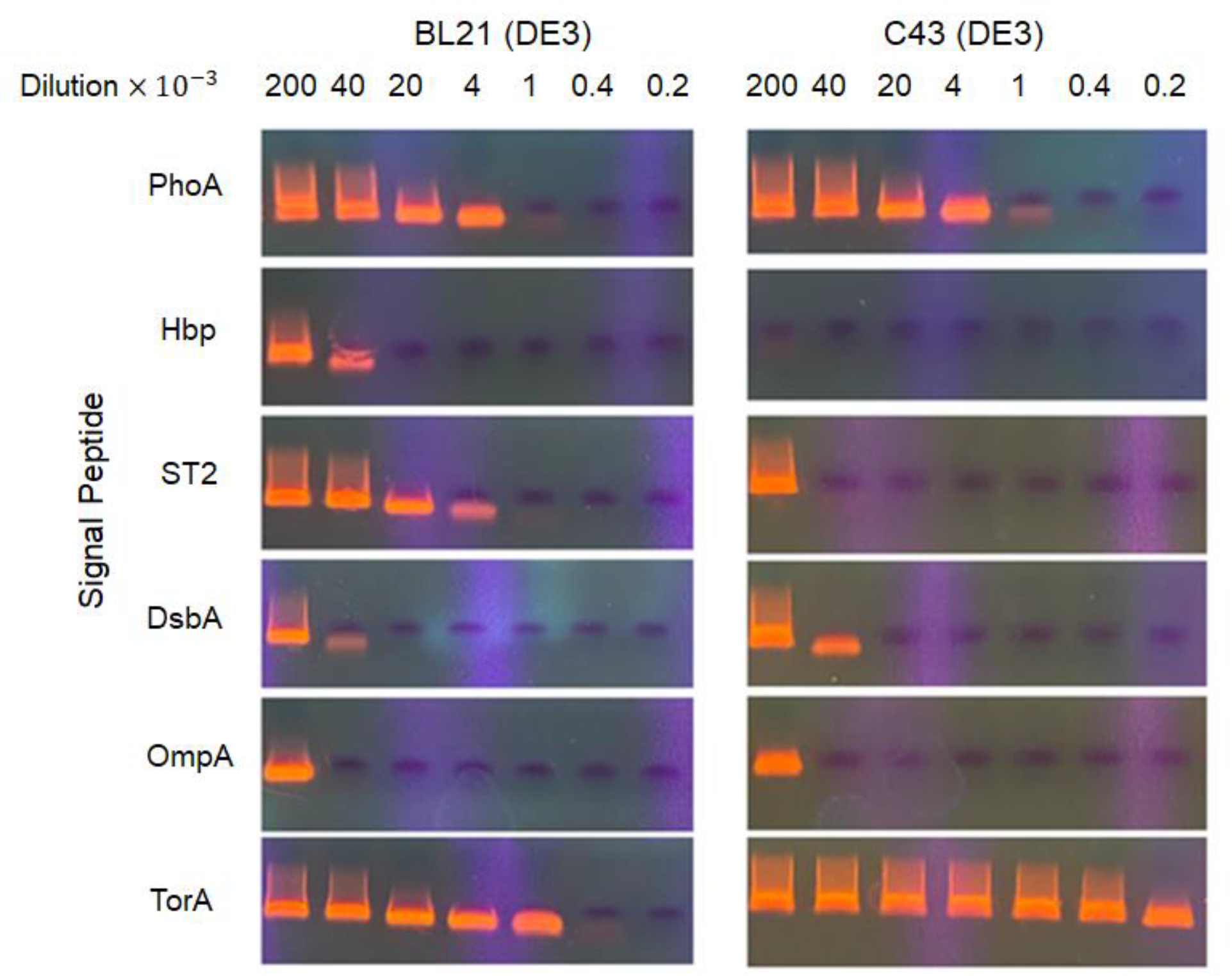
1% agarose gels illustrating nuclease activity of E. coli cell lysates in which SmEn was secreted using different signal peptides in E. coli BL21 (DE3) (left) or C43 (DE3) (right).

Using C43 (DE3) instead of the conventional BL21 (DE3) host was most beneficial with the ST2 signal peptide resulting in ∼40 times higher SmEn activity. Interestingly, minimal or no activity increase was gained by switching to the C43 (DE3) host when using the PhoA, DsbA, and OmpA signal peptides.

To better understand signal peptide influences, the most probable RNA secondary structures for the translation initiation region of the mRNAs were determined. Gibbs free energy for structure stabilities were determined for the sequence composed of the 30 nucleotides upstream of the translation initiation codon and the initial 40 downstream nucleotides of the structural gene encoding the different signal sequences. Previous studies indicate that stable mRNA secondary structure in this region can slow translation initiation [41-43]. As shown in Table S2, mRNAs encoding the PhoA, OmpA, and TorA signal peptides are estimated to have the highest probability of strong secondary structure according to their Gibbs free energies as opposed to sequences for ST2, DsbA, and Hbp signal peptides with higher values. Evidently, secretion with the PhoA and OmpA signal peptides is not improved using C43 (DE3) because the expression rates are already restrained by mRNA secondary structures that impede translation initiation. Lower SmEn activity accumulation using the TorA signal sequence could be caused either by mRNA secondary structure or a lower translocation rate supported by the Tat system or potentially both.

### SmEn purification and testing nuclease activity

The purification protocol was developed using cell lysates of C43 (DE3) cells expressing the SmEn gene with the Hbp signal peptide. Figure 3 shows a SDS-PAGE gel of Ni-NTA affinity purification fractions.

**Figure 3.**
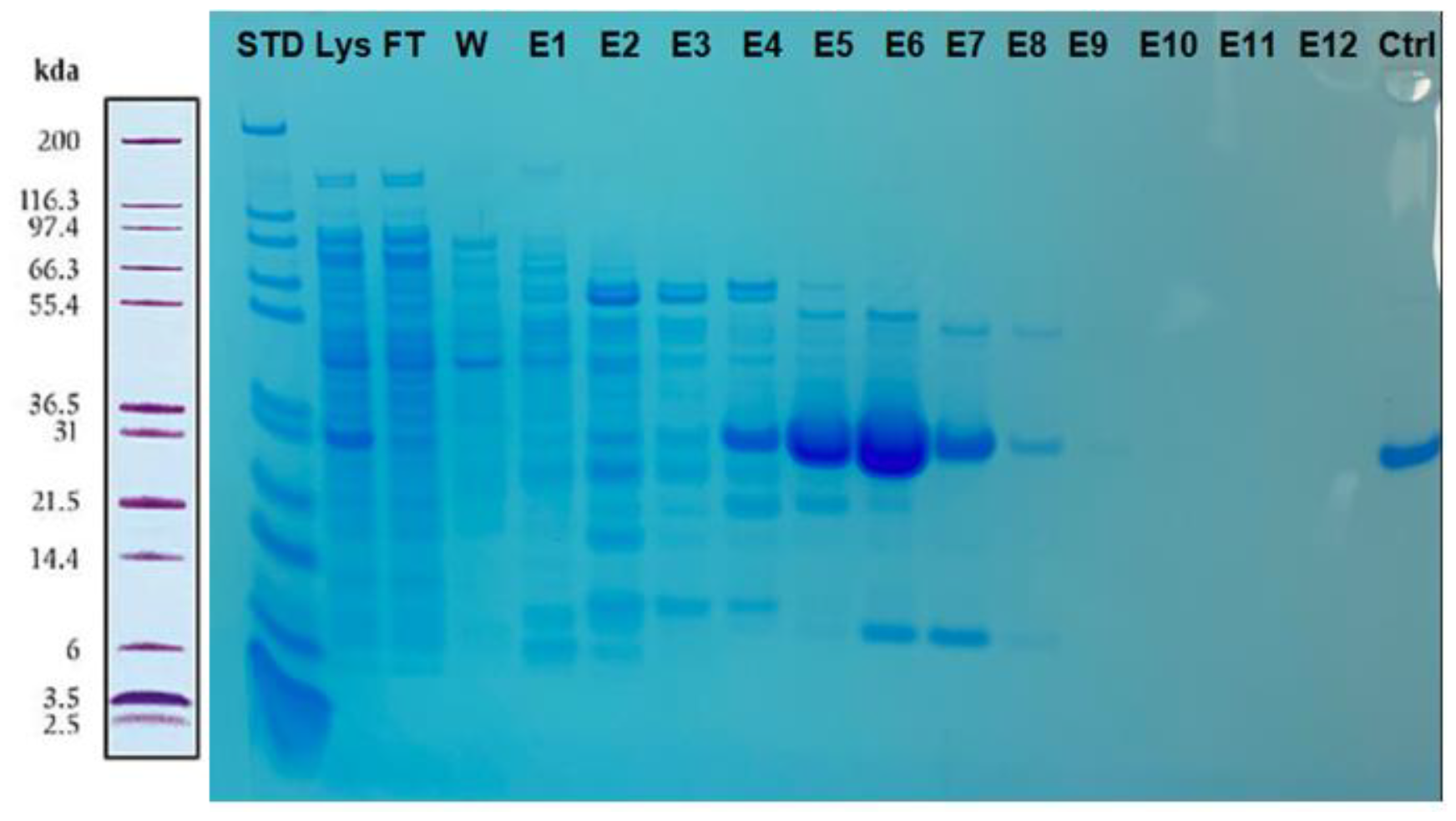
SDS-PAGE visualizing SmEn purification by Ni-NTA resin. Fractions E5-E7 were combined and dialyzed. FT= Flow Through, W= Wash, E= Elution.

As mentioned previously, lysate before purification as well as flow through and wash were diluted 15 times before loading onto the protein gel. Elution fractions 5,6, and 7, eluted with 150-200 mM imidazole, were combined and dialyzed against Z buffer three times to remove imidazole. About 54 mg of SmEn was prepared from a 1 L culture as measured by the DC protein assay.

To fully validate the new process, the specific activity of SmEn translocated by the PhoA signal peptide was compared to that of the commercial product from Millipore (Cat# 101654). Figure 4A shows that the secreted E. coli and commercial sample of SmEn have significant activity. For complete degradation of 0.74 µg/µl salmon sperm DNA at 37 °C in 30 minutes 200 ng/ml of pure SmEn is needed as indicated by both in-house and by Millipore SmEn, indicating approximately equivalent specific activity, that is full activation of the E. coli secreted enzyme.

**Figure 4.**
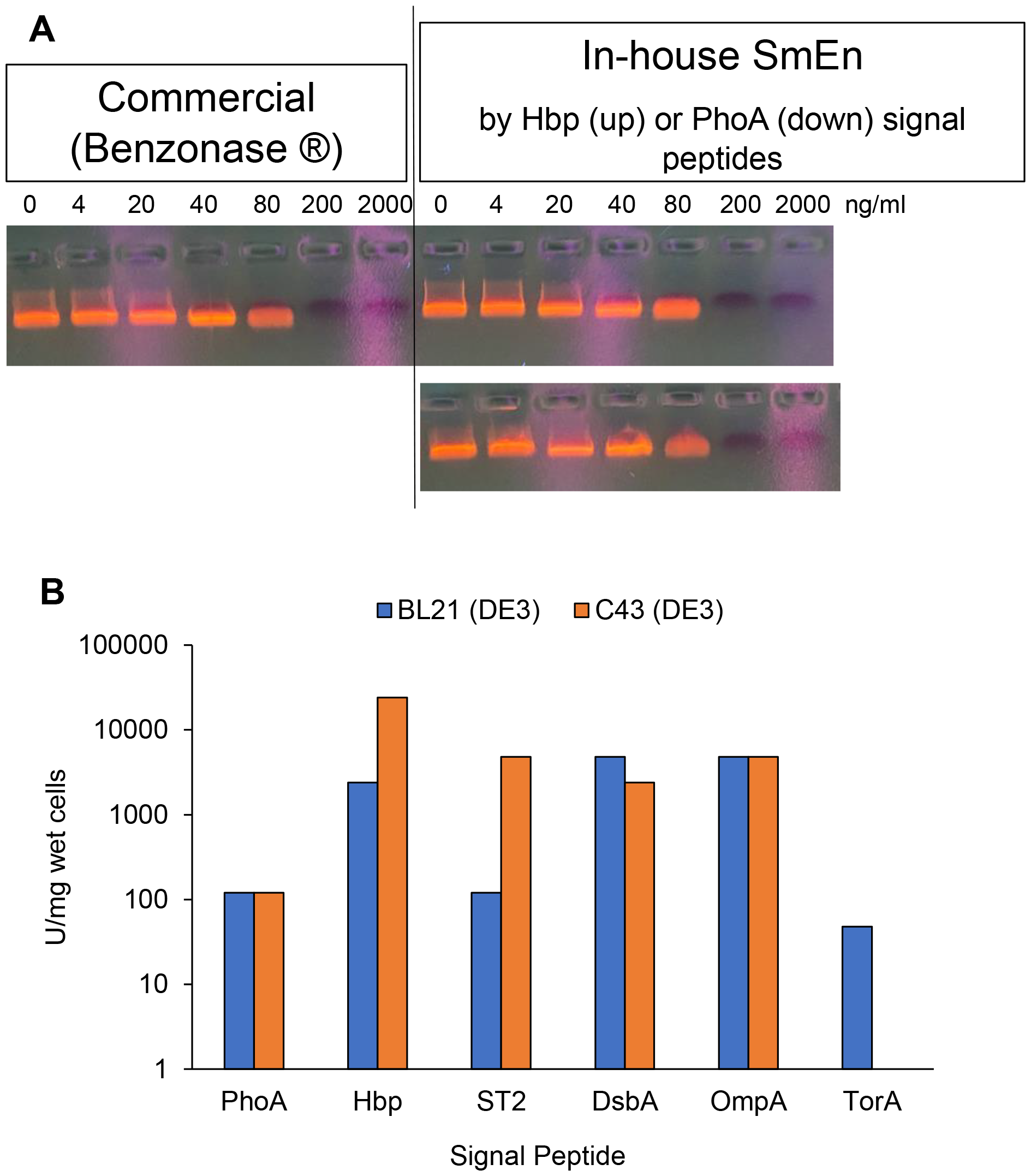
A) Nuclease activity of SmEn prepared from C43 (DE3) cells translocating SmEn using the PhoA signal peptide at 26 °C (left) versus commercial SmEn (Millipore). B) Estimated SmEn activity accumulation per gr wet cells for SmEn translocation and activation in BL21 (DE3) (blue bars) or C43 (DE3) (orange bars) using different signal peptides.

According to Figure 4A, 200 ng/ml SmEn is needed to fully digest 0.74 µg/µl salmon sperm DNA at 37 °C in 30 minutes which is defined as 1 U. This calibration allows estimation of the active SmEn yields based on the results shown in Figure 2. These are indicated in Figure 4B on a logarithmic scale with the highest SmEn activity accumulation of about 24000 U/mg wet cells (equivalent to 240 mg SmEn/g wet cells) when using the Hbp signal sequence and C43 (DE3).

## Conclusion

Potential cytoplasmic toxicity and disulfide bond formation associated with active SmEn recombinant expression in E. coli motivated SmEn’s translocation to the periplasmic space. We explored the impacts of post-induction culture incubation temperature, IPTG induction level, E. coli host strain, different signal sequences, and the RNA secondary structure at the translation initiation region for SmEn translocation. This work used a new rapid SmEn activity accumulation assay to avoid the need for protein purification and buffer exchange. This assay significantly reduces the assessment time to less than 1.5 hours from cell harvest to specific activity determination.

Our findings suggest that higher growth temperatures should be avoided, as incubation at 37 °C most likely reduced overall protein production yields because of the rapid synthesis of the polypeptide chain overwhelmed the translocation machineries [27, 34, 37]. Incubation at room temperature resulted in slower protein production and translocation preventing common issues associated with rapid membrane protein production, including cell stress responses, inclusion body formation, and even growth arrest [18].

Among the tested signal sequences, the TAT system resulted in the lowest SmEn accumulation as expected considering post-translational translocation using the Tat pathway. On the other hand, the Hbp and OmpA signal sequences provided the highest SmEn activity accumulation, regardless of the E. coli strain used. We also observed that the C43 (DE3) E. coli host with weakened transcription improved SmEn translocation more than 40 times in comparison to the BL21 (DE3) stain for the ST2 signal peptide. However, this improvement was minimal or absent for other signal sequences. This discrepancy may stem from the relative lack of RNA secondary structure at the TIR for the ST2 signal sequence where absence of a stable secondary structure necessitates slower transcription and subsequently, translation and translocation.

Our analysis of mRNA secondary structures at the TIR indicated that while PhoA and OpmA had comparable overall Gibbs free energies, the PhoA signal sequence has a much stronger secondary structure in the TIR compared to the OmpA signal sequence. This suggests that Gibbs free energy for mRNA secondary structures should focus more specifically on the TIR. Silent mutation of signal sequences to adjust mRNA secondary structures in this region could improve secreted protein production and translocation. Overall, our study sheds light on crucial factors influencing SmEn translocation and activation providing valuable insights for optimizing protein production and translocation processes in bacterial systems.

