## Supplementary Material for "Secretion and Periplasmic Activation of a Potent Endonuclease in E. coli"

**Address:** 443 Via Ortega, 093, Stanford, CA 94305

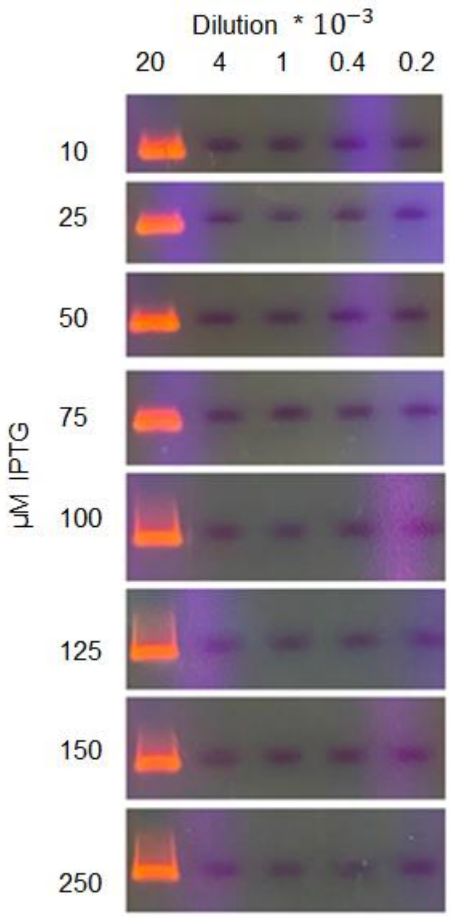


Figure S1. Lack of an IPTG titration effect on SmEn activity accumulation using the Hbp signal peptide in C43 (DE3) cells.

Table S1. Final $OD_{600}$ for SmEn production after overnight incubation at room temperature by BL21 (DE3) or C43 (DE3) host cells.

| | *BL21 (DE3)* | *C43 (DE3)* |
|---|---|---|
| ST2 | 4.12 | 3.01 |
| DsbA | 4.55 | 3.22 |
| TorA | 4.04 | 3.53 |
| OmpA | 4.15 | 3.38 |
| Hbp | 4.14 | 3.77 |
| PhoA | 3.59 | 3.86 |
| No SP | 3.77 | Not tested |

Table S2. RNA secondary structure analysis and amino acid composition of different signal peptides

| | **RNA secondary structure analysis** | | | | **Amino acid compositions (%)** | | | |
|---|---|---|---|---|---|---|---|---|
| **Signal Peptide** | **Energy (kcal/mol)** | **$2^{nd}$ structure before AUG** | **Translation initiation** | **$2^{nd}$ structure after AUG** | **Hydrophobic** | **Acidic** | **Basic** | **Neutral** |
| **PhoA** | -9.4 | - | Stem (>80%) | Stem-loop close-(>95%) | 68.18 | 0 | 9.09 | 22.73 |
| **Hbp** | -3.8 | Short Loop, close (>90%) | Stem (>50%) | Stem-Loop, far (>80%) | 55.77 | 1.92 | 17.31 | 25 |
| **ST2** | -3.3 | Loop (>70%) | Stem (50%>) | Stem-Loop, relatively close (>80%) | 65.22 | 0 | 8.7 | 26.09 |
| **DsbA** | -3.3 | - | Loop (>90%) | Loop, close (>90%) | 73.68 | 0 | 10.53 | 15.79 |
| **OmpA** | -8.2 | Short loop, close (>60%) | Stem (>50%) | Stem-Loop -far (>99%) | 71.43 | 0 | 9.52 | 19.05 |
| **TorA** | -5.9 | Short loop-close (>90%) | Stem (50%>) | Stem Loop -far (>80%) | 48.72 | 2.56 | 12.82 | 35.9 |

Sm Endonuclease (SmEn) sequence. The SmEn sequence (PDB ID: 1G8T) without signal sequence is underlined and highlighted green. The RBS sequence is highlighted in blue. The T7 promoter and T7 terminator sequences are in red. BmtI and BlpI restriction sites are italic.

TAATACGACTCACTATAGGGGAATTGTGAGCGGATAACAATTCCCCTCTAGAAATAATTTTGTTTAACTTTAAG
AAGGAGATATACATATG*GCTAGC*GATACCCTGGAAAGCATTGATAATTGTGCAGTTGGTTGTCCGACCGGT
GGTAGCAGCAATGTTAGCATTGTTCGTCATGCATATACCCTGAATAATAACAGCACCACCAAATTTGCCAAT
TGGGTTGCATATCACATCACCAAAGATACACCGGCAAGCGGTAAAACCCGTAATTGGAAAACCGATCCG
GCACTGAATCCGGCAGATACACTGGCACCGGCAGATTATACCGGTGCAAATGCAGCACTGAAAGTTGAT
CGTGGTCATCAGGCACCGCTGGCAAGCCTGGCAGGCGTTAGCGATTGGGAAAGCCTGAATTATCTGAG
CAATATTACACCGCAGAAAAGCGATCTGAATCAAGGTGCATGGGCACGTCTGGAAGATCAAGAACGTAAA
CTGATTGATCGTGCAGATATCAGCAGCGTTTATACCGTTACCGGTCCGCTGTATGAACGTGATATGGGTAAA
CTGCCTGGTACACAGAAAGCACATACCATTCCGAGCGCATATTGGAAAGTGATCTTTATTAACAATAGCCC
TGCCGTGAATCACTATGCAGCATTTCTGTTTGATCAGAATACCCCGAAAGGTGCAGATTTTTGTCAGTTTCGT
GTTACCGTGGATGAAATTGAAAAACGTACCGGTCTGATTATTTGGGCTGGCCTGCCGGATGATGTTCAGGC
GAGCCTGAAAAGCAAACCGGGTGTGCTGCCGGAACTGATGGGTTGTAAAAATGGTAGCGGTAGCCATCA
CCATCATCATCATTAA*GCTGAGC*AATAACTAGCATAACCCCTTGGGGCCTCTAAACGGGTCTTGAGGGGTT
TTTTGCTGAAAGGAGGAACTATATCCGGAT

DNA sequences for signal peptides.

| Signal sequence | DNA sequence |
|---|---|
| PhoA | ATGGTTAAACAATCTACCATTGCTCTGGCACTGCTGCCGCTGCTGTTTACTCCGGTTACTAAAGCGC |
| Hbp | ATGAACCGTATCTACTCCCTGCGTTACTCCGCTGTAGCTCGTGGTTTCATCGCGGTTTCTGAATTCGCGCGCAAATGCGTCCATAAAAGCGTGCGCCGCCTGTGCTTCCCGGTTCTGCTGCTGATCCCAGTGCTGTTTTCTGCTGGCTCTCTGGCC |
| ST2 | ATGAAAAAGAATATCGCATTTCTTCTTGCATCTATGTTCGTTTTTTCTATTGCTACAAATGCCTATGCA |
| DsbA | ATGAAAAAGATCTGGCTGGCACTGGCTGGTCTGGTTCTGGCTTTTTCCGCTTCTGCA |
| OmpA | ATGAAAAAAACTGCCATCGCTATCGCGGTGGCTCTGGCTGGCTTCGCAACCGTAGCGCAGGCT |
| TorA | ATGAACAACAACGACCTGTTTCAGGCGTCTCGTCGTCGTTTCCTGGCGCAACTGGGTGGCCTGACCGTTGCGGGCATGCTGGGCCCGTCCCTGCTGACCCCGCGCCGCGCAACTGCG |
